# Heterologous boost with mRNA vaccines against SARS-CoV-2 Delta/Omicron variants following an inactivated whole-virus vaccine

**DOI:** 10.1101/2022.09.06.506714

**Authors:** Changrui Lu, Yuntao Zhang, Xiaohu Liu, Fujun Hou, Rujie Cai, Zhibin Yu, Fei Liu, Guohuan Yang, Jun Ding, Jiang Xu, Xianwu Hua, Xinping Pan, Lianxiao Liu, Kang Lin, Zejun Wang, Xinguo Li, Jia Lu, Qiu Zhang, Yuwei Li, Chunxia Hu, Huifeng Fan, Xiaoke Liu, Hui Wang, Rui Jia, Fangjingwei Xu, Xuewei Wang, Hongwei Huang, Ronghua Zhao, Jing Li, Hang Cheng, William Jia, Xiaoming Yang

**Affiliations:** China National Biological Group-Virogin Biotech (Shanghai) Ltd (CNBG-Virogin); China National Biotec Group (CNBG); Virogin Biotech (Shanghai) Ltd (Virogin); Wuhan Institute of Biological Products Co., LTD (WIBP); Beijing institute of biological products Co., LTD; Shuimu BioSciences Ltd.

## Abstract

The coronavirus SARS-CoV-2 has mutated quickly and caused significant global damage. This study characterizes two mRNA vaccines ZSVG-02 (Delta) and ZSVG-02-O (Omicron BA.1), and associating heterologous prime-boost strategy following the prime of a most widely administrated inactivated whole-virus vaccine (BBIBP-CorV). The ZSVG-02-O induces neutralizing antibodies that effectively cross-react with Omicron subvariants following an order of BA.1>BA.2>BA.4/5. In naïve animals, ZSVG-02 or ZSVG-02-O induce humoral responses skewed to the vaccine’s targeting strains, but cellular immune responses cross-react to all variants of concern (VOCs) tested. Following heterologous prime-boost regimes, animals present comparable neutralizing antibody levels and superior protection across all VOCs. Single-boost only generated ancestral and omicron dual-responsive antibodies, probably by “recall” and “reshape” the prime immunity. New Omicron-specific antibody populations, however, appeared only following the second boost with ZSVG-02-O. Overall, our results support a heterologous boost with ZSVG-02-O, providing the best protection against current VOCs in inactivated virus vaccine– primed populations.

## INTRODUCTION

The coronavirus SARS-CoV-2 and its developing variants spread quickly across borders, causing hundreds of millions of infected cases and several millions of deaths worldwide as of 2021 ^1^.

Currently our most potent countermeasure, COVID-19 vaccines have a number of platforms including inactivated whole-virus ^2^, subunit vaccines ^3^, lipid nanoparticles (LNP) mRNA ^4, 5^, adenoviral vectors expressing viral Spike protein ^6^ and various recombinant S protein vaccines and DNA vaccines ^7^. While mRNA vaccines induce higher antibody titers, inactivated whole-virus vaccines have had many years of manufacture and clinical experience. Moreover, inactivated whole-virus vaccines may induce a broader breadth of immunity, specifically for viral structure proteins other than just the S protein.

While the current vaccines have proven very effective against the original COVID-19 (SARS-CoV-2 ancestral strain), emerging new variants of concern (VOCs) have quickly reduced their efficacy due to mutations that escape antibody neutralization ^8-11^. Recent variants, including BA.1 ^12^, BA.2 ^13^, BA.3 (B.1.1.529.3) ^14^ and BA4/5 ^15^, can resist neutralizing antibodies induced by either convalescence or currently approved COVID-19 vaccines ^16-20^. While booster shots of BNT162b2 or mRNA-1273 effectively prevent severe symptoms, breakthrough infections with the Omicron variants in people have been widely reported ^21^. Therefore, updated booster vaccines and associated immunization schemes against new VOCs remain urgently necessary.

Given the advantages of inactivated whole-virus vaccines in immunity breadth and a vast population have been primed with them in China and worldwide, an effective boost strategy against current Omicron variants requires extensive studies ^22^. Recent studies show that two doses of the inactivated whole-virus vaccine, followed by one mRNA (BNT162b2 or mRNA-1273) as the booster, can prominently increase the humoral and cellular immune responses to COVID variants, but less effective for Omicron variant^23 24^. Other studies reported that a viral vector vaccine primer plus an mRNA booster exhibited a high level of cross-neutralizing activity for the ancestral, Beta, Delta, and Omicron strains ^25, 26^. Clinical studies of heterologous boost with mRNA and adenovirus vaccines against the ancestral SARS-CoV-2 following two doses of the BBIBP-CorV vaccine have also been conducted ^27^. While the heterologous boost vaccination was safe, the inhibition for omicron variant measured less than 70% by a surrogate neutralization test ^28^. Inactivated whole virus vaccines of ancestral SARS-CoV-2, such as Sinopharm (BBIBP-CorV) and Sinovac (CoronaVac), have reached worldwide popularity and helped to mitigate the pandemic in many countries, especially in less developed areas ^29, 30^. It is important to demonstrate that a heterologous boosting with a mRNA vaccine designed for omicron variants following the above inactivated virus vaccine can be effectively against Omicron variants and furthermore, can broaden the immunity for multiple variants of SARS-CoV-2.

Here we report two new SARS-CoV-2 mRNA vaccines, ZSVG-02 and ZSVG-02-O, against Delta B.1.617.2 variant and Omicron B.1.1.529.1 (BA.1 subvariant), respectively. Both induce high neutralizing antibody levels against the variants in mouse models. In addition, the ZSVG-02-O-induced neutralizing antibodies can cross react with Omicron subvariant BA.2 and BA.4/5, albeit with reduced activity. Serving as a boost to BBIBP-CorV ^27^, both vaccines induce additional protection across all VOCs tested in an ACE2-expressing mouse model. Moreover, our results demonstrate that a heterologous boost strategy with ZSVG-02-O following BBIBP-CorV priming produces superior protective immunity over homologous boosts. The heterologous prime-boost strategy with BBIBP-CorV and ZSVG-02-O takes advantages of both the primer and booster vaccines that the mRNA vaccines targeting the new VOCs and also recalling a broad breadth of immunity against SARS-Cov-2 built by the previously primed BBIBP-CorV.

## RESULTS

### ZSVG-02- and ZSVG-02-O-encoded active Spike proteins and structural analysis with cryo-EM

In order to confirm structures of the Delta and Omicron Spike proteins expressed by ZSVG-02 and ZSVG-02-O vaccines, we first tested their expression in transfected cells. The mRNA-LNPs were prepared and characterized following a classic protocol (Figure S1A-D). Naked or LNP-encapsulated, ZSVG-02 and ZSVG-02-O expressed their corresponding S protein in transfected HEK293 cells (Figure 1A, S1E). Specifically, the LNP ensured nearly 100% transfection efficiency of both ZSVG-02 and ZSVG-02-O in cells, measured by flow cytometry using a labeled angiotensin-converting enzyme 2 (ACE2) protein (Figure 1B). The resulting S-protein antigens expressed by the transfected cells can also bind to receptor-binding domain (RBD)-specific antibodies against other variants (Data not shown). Antibodies against RBD can completely displace ACE2 binding to the ZSVG-02 transfected cells, showing that it competes with ACE2 for Spike binding (Figure S1F).

**Figure1.**
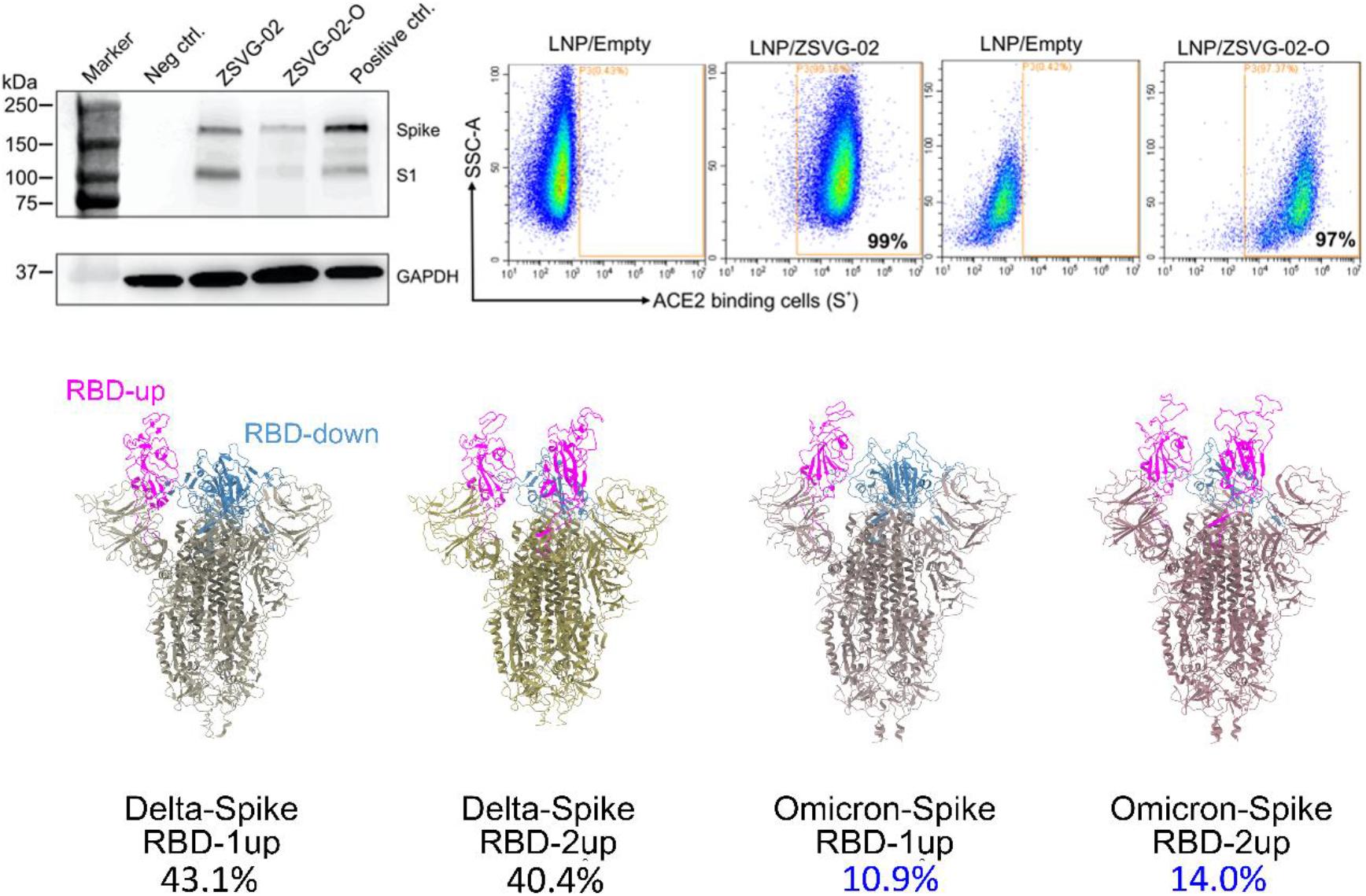
Naked or LNP-encapsuled mRNA transfected to HEK293T/17 cells show specific spike protein expression and reduced immunogenicity. The 3D EM maps and 3D models for different Delta and Omicron Spike proteins produced from ZSVG-02 and ZSVG-02-O mRNA. (A). Detection of SARS-CoV-2 spike proteins expressed by naked mRNA of either ZSVG-02 or ZSVG-02-O in cells lysate via Western Blot. Cell lysates of mock (empty lipofectamine 3000) and pcDNA3.1(-) plasmid expressing S-protein transfections as negative and positive controls, respectively. n=4, (B). Representative flow plots from 4 independent experiments showing the expression of LNP-encapsulated ZSVG-02 or ZSVG-02-O on cell surface by binding with 2.5ug/ml ACE2 protein. The threshold was set according to the empty LNP transfected cells. (C) 3D models built from EM maps. Delta spike active state with 2 up RBD is colored with corn khaki. Omicron spike active state with 1 up RBD is colored with misty rose. Omicron spike active state with 2 up RBD is colored with misty pink. The up state RBD is highlighted with magenta. The down state RBD is highlighted with steel blue. Numbers underneath each model indicate the percentage of each conformation within total particles after 2D classification for Delta and Omicron respectively.

Next, we used cryogenic electron microscopy (Cryo-EM) and single particle analysis (SPA) methods to determine the 3D structures of the two variants. We determined the overall resolution of Delta Spike protein open state with 1 receptor-binding domain (RBD) up at 2.64 Å, 2 RBD up at 2.77 Å, and Omicron Spike protein open states with 1 RBD up at 3.02 Å, and 2 RBD up at 3.02 Å respectively (Figure S1I-J). We have built the 3D models for all four cryo-EM maps that covered most domains (Figure 1C). Our 3D Heterogeneous Refinement shows that amongst all 2D particles for the Delta Spike protein, 43.1% belongs to the 1up-2down conformation, while 40.4% belongs to the 2up-1down conformation. The Omicron Spike showed 10.9% 1up-2down conformation and 14% 2up-1down conformation (Figure 1C). Compared to the Delta Spike, the Omicron counterpart shows greater flexibility with more than 60% particles belonging to the “middle” RBD conformation, somewhere between up and down (data not shown). Figure S1K shows the local resolution for all the models built. The Omicron RBD domains show lowest resolution compared to Delta counterparts, indicating more structural dynamic.

### ZSVG-02-O elicits strong humoral and cellular immune response against Omicron subvariants

To determine the ability of the mRNA vaccines to induce neutralizing antibodies (NAbs) against current SARS-CoV-2 VOCs, BALB/c mice received two doses of ZSVG-02-O over a two-week interval (Figure 2). The sera were collected 7 days following the second immunization to measure neutralizing antibodies against Omicron (BA.1) live virus. Figure 2A shows that ZSVG-02-O induces high levels of anti-Omicron neutralizing antibodies, while levels against ancestral strain and Delta strains are moderate for the middle and high doses and barely detectable for the lowest dose. This result proves variant specificity of ZSVG-02-O-induced immune response directly targeting the Omicron variant. Parallel experiments with the Delta-specific vaccine ZSVG-02 showed similar results with the most neutralizing antibodies against Delta followed by that for the ancestral strain. However, the antibody levels for Omicron variant BA.1 drop by 60- and 16-fold compared to Delta for the two doses tested (Figure S2).

**Figure 2:**
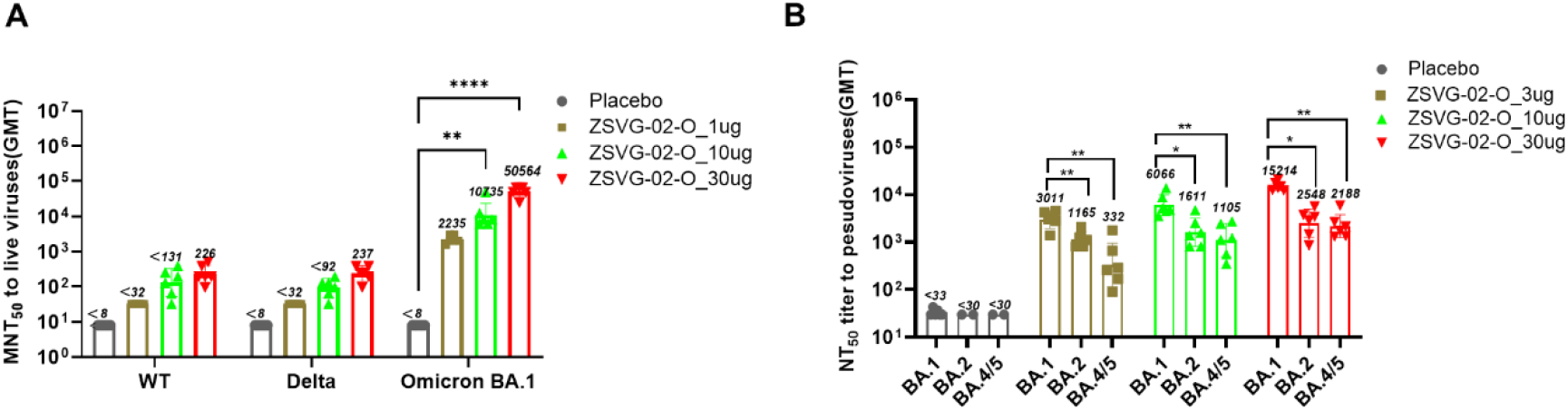
Serological evaluation of ZSVG-02-O immunization in mice received 2 doses of ZSVG-02-O with a 14-day interval. (A) GMT of neutralizing antibodies against live SAR-CoV-2 variants one week after the second immunization. (B) GMT of neutralizing antibodies against Omicron BA.1, BA.2 and BA.4/5 pseudoviruses two weeks after the second immunization. WT, ancestral strain, * p<0.05, ** p<0.01, **** p<0.0001

To investigate ZSVG-02-O-induced humoral immunity across the current Omicron subvariants, the sera of ZSVG-02-O-vaccinated animals were collected fourteen days following the second dose and NAb levels were determined with pseudoviruses of three Omicron subvariants (BA.1, BA.2, BA.4/5) (Figure 2B). Figure 2B shows that levels of NAbs elicited by ZSVG-02-O reduced by 2.6-, 3.8- and 6.0-fold for BA.2 and 16.5-, 24.3- and 6.2-fold for BA.4/5, respectively at 3 µg, 10 µg, and 30 µg doses.

T cell immune response in animals received two doses of ZSVG-02-O (3 µg, 10 µg, and 30 µg) with a 14-day interval was determined. Splenocytes were collected and analyzed 4 weeks after the second vaccination. Spike protein-specific cellular immunity was evaluated by intracellular staining of the cytokines induced by incubating the splenocytes with the full-length S protein of different variants. Figures 3A-H show that ZSVG-02-O-elicited antigen-specific response in both CD4+ and CD8+ T cells, indicated by the elevated percentages of IFNγ+/CD69+ T cells following the treatment with ancestral, Delta, or Omicron BA.1 Spike proteins, respectively. Parallel ELISpot assay for interleukin (IL)-2 expression in splenocytes further confirmed a strong immune response against the Omicron (BA.1) variant by ZSVG-02-O-immunized mice stimulated with Omicron Spike protein (Figure 3I). Meanwhile, IL-5 was not significantly elevated across all samples (Figure 3J), suggesting that ZSVG-02-O mainly induced a Spike-specific Th1 immune response rather than Th2. We found no T cell immunity against the nucleocapsid protein of the ancestral virus (Figure 3D and H) in ZSVG-02-O-vaccinated animals. Similarly, the Delta-specific vaccine ZSVG-02 also induced significantly elevated cellular immune responses against Spike proteins of the ancestral, Delta, and Omicron variants but not the ancestral nucleocapsid protein. (Figure S3).

**Figure 3:**
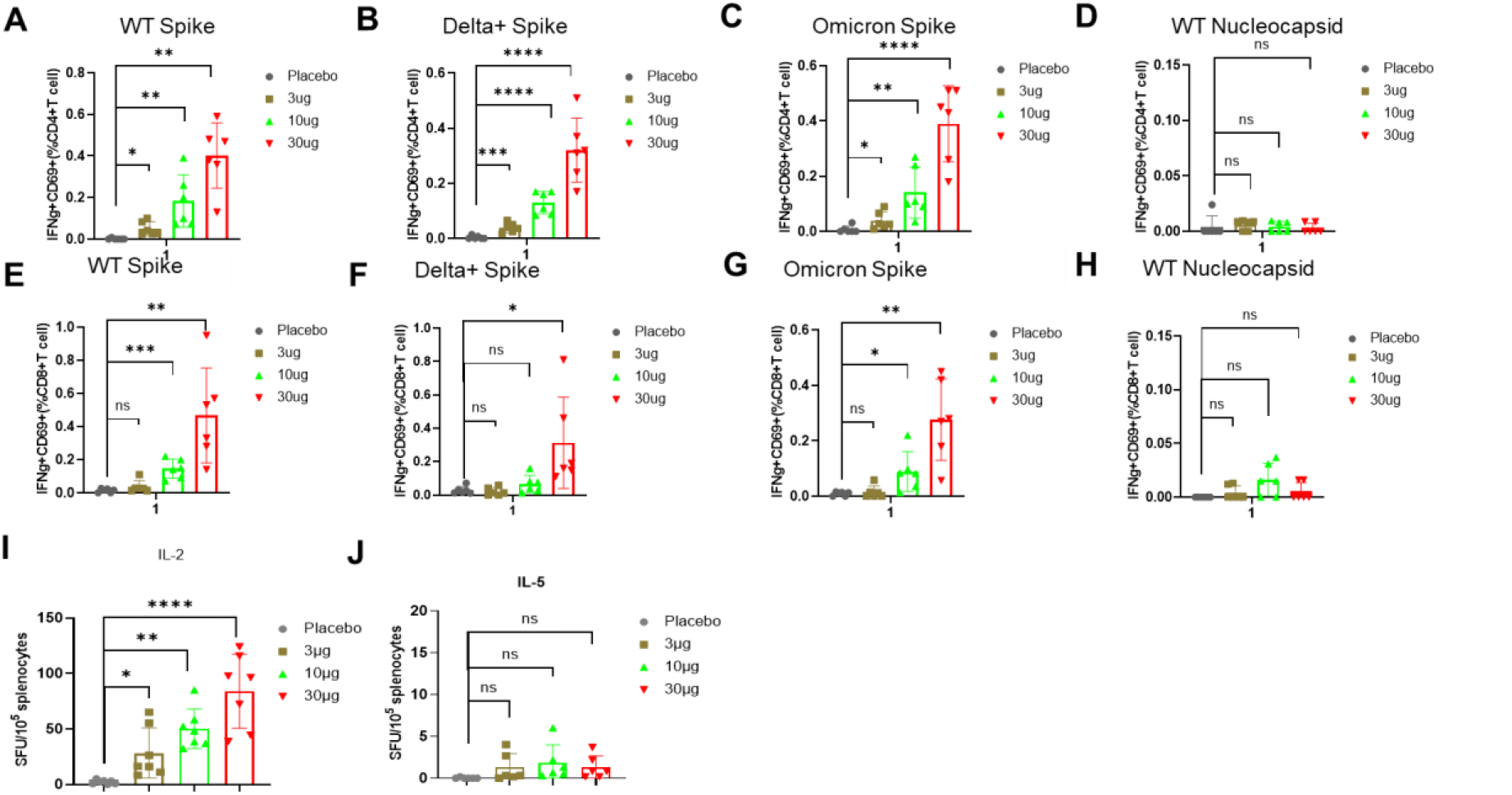
SARS-CoV-2 Spike-specific T-cell immune response in ZSVG-02-O-vaccinated mice received 2 doses of ZSVG-02-O with a 14-day interval. Splenocytes were collected 4 weeks post second immunization and stimulated with the indicated proteins of different variants for 48 hours. (A)-(H) The proportions of IFNγ+/CD69+ CD4+ and CD8+ T cells from ZSVG-02-O immunized mice stimulated with spike proteins or WT nucleocapsid protein. (I)-(J) ELISpot assay for IL-2 (I) and IL-5 (J) in splenocytes stimulated with Omicron Spike protein. Data are shown as mean ± SD (* p<0.05, ** p<0.01, *** p<0.001, ****p < 0.0001). WT, ancestral strain.

Together, these results show that ZSVG-02 and ZSVG-02-O induce strong protective immune responses against SARS-CoV-2 variants at both humoral and cellular levels.

### Heterologous boost of ZSVG-02-O following two-dose BBIBP-CorV primer induced strong protective immune response

Since a large proportion of the world had received inactivated whole-virus vaccines of ancestral strain, we further investigated whether boosting with the Omicron-specific ZSVG-02-O mRNA vaccines following an inactivated whole-virus vaccine could induce greater protective immune response against various VOCs. For that, BALB/c mice were first Intramuscularly (IM) immunized with two doses of BBIBP-CorV followed by one dose of ZSVG-02-O as a heterologous boost (Figure 4A). Figure 4B-C show the NAb levels determined with live viruses of the three strains following the two prime-boost regimes tested. Homologous BBIBP-CORV boost can produce adequate NAb levels against the ancestral strain and Delta variant but not for Omicron BA.1. A single ZSVG-02-O boost, however, can induce NAbs of comparable levels across all three strains. Fourteen days post ZSVG-02-O boost further increased the NAb levels compared to 7 days post immunization, especially against Omicron BA.1. Notably, the NAb levels induced by the single ZSVG-02-O boost following two-dose BBIBP-CorV prime show no obvious dose dependency, and the antibodies induced show the highest level against the ancestral strain.

**Figure 4.**
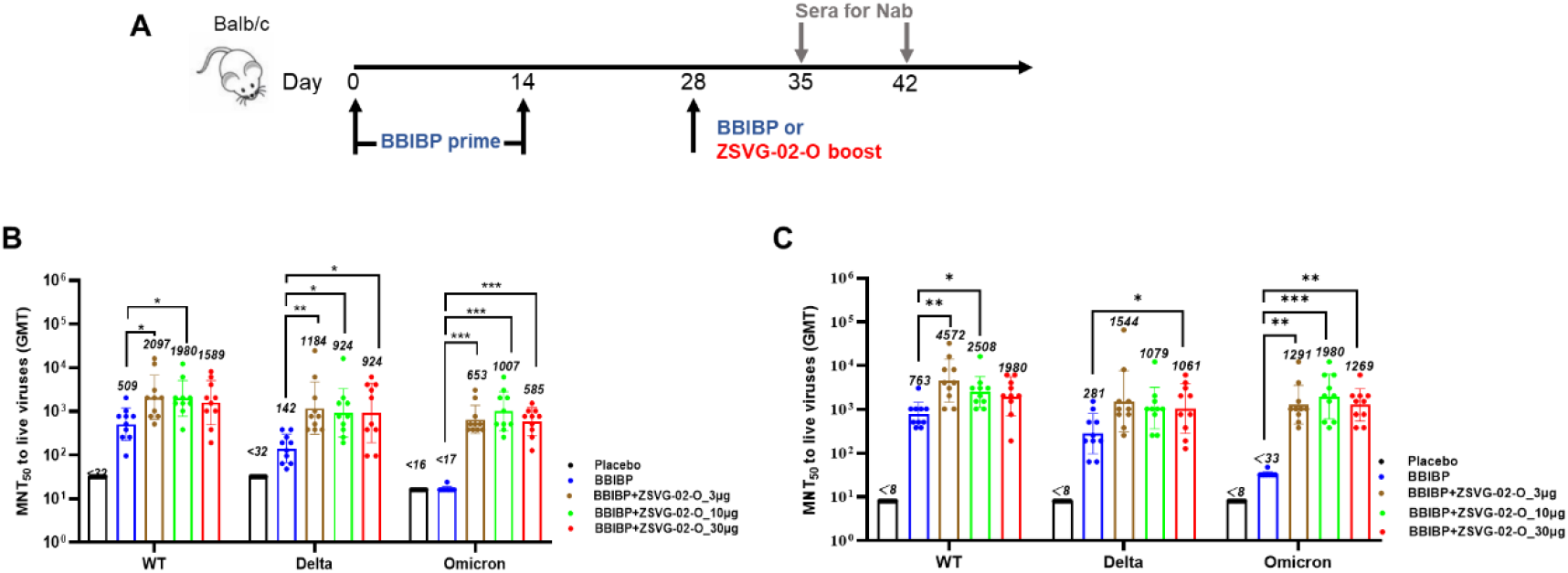
Serological evaluation of single-dose ZSVG-02-O boost following 2-dose immunization of inactivated vaccine in mice. (A) Schematic diagram of immunization regimens. (B) Neutralizing antibody titers (GMT) measured in sera of mice seven days post vaccination (Day 35) with the ZSVG-02-O boost. (C) Neutralizing antibody titers fourteen days post vaccination (Day 42) with the ZSVG-02-O boost. WT, ancestral strain.

### ZSVG-02-O heterologous boost protected mice from challenges by Delta and Omicron BA.1 viruses

Next, we tested the protection by the above ZSVG-02-O heterologous boost on ACE2-expressing transgenic mice, challenged with Delta or Omicron variant live viruses. The immunization scheme and experiment timing are shown in Figure 5A. Fourteen days following the boost, the animals were challenged intranasally with the viruses at a dose of 200 CCID_50_. In the saline control group, we observed a significant body weight loss starting from Day 4 (Figure 5B) after Delta or Omicron variant challenges. Four out of four mice died 6 days post challenge in the Delta group. Two out of four mice died and the remaining two were severely sick 8 days post infection with the Omicron variant (Figure 5C). No animal died or lost body weight in the ZSVG-02-O heterologous boost group 8 days post infection with Omicron. The 1 µg and 10 µg groups boosted with ZSVG-02-O showed moderate body weight loss in Delta variant-challenged animals, while the 30 µg group remained healthy. Furthermore, virus loads of both VOCs in heterologously vaccinated animals fell significantly below the controls, measured by qRT-PCR tests on turbinate and lung tissue (Figure 5E,F). By contrast, the BBIBP-CORV homologous boost group showed notable body weight loss after Delta variant challenge with no death observed. This group also survived Omicron challenge with no measurable body weight loss and showed reduced virus loads in tissues compared to the saline group (Figure 5C-F). Overall, for the BBIBP-CorV primed mouse, ZSVG-02-O heterologous boost offered 100% protection against both Delta and Omicron virus, superior to a third homologous BBIBP-CorV boost.

**Figure 5:**
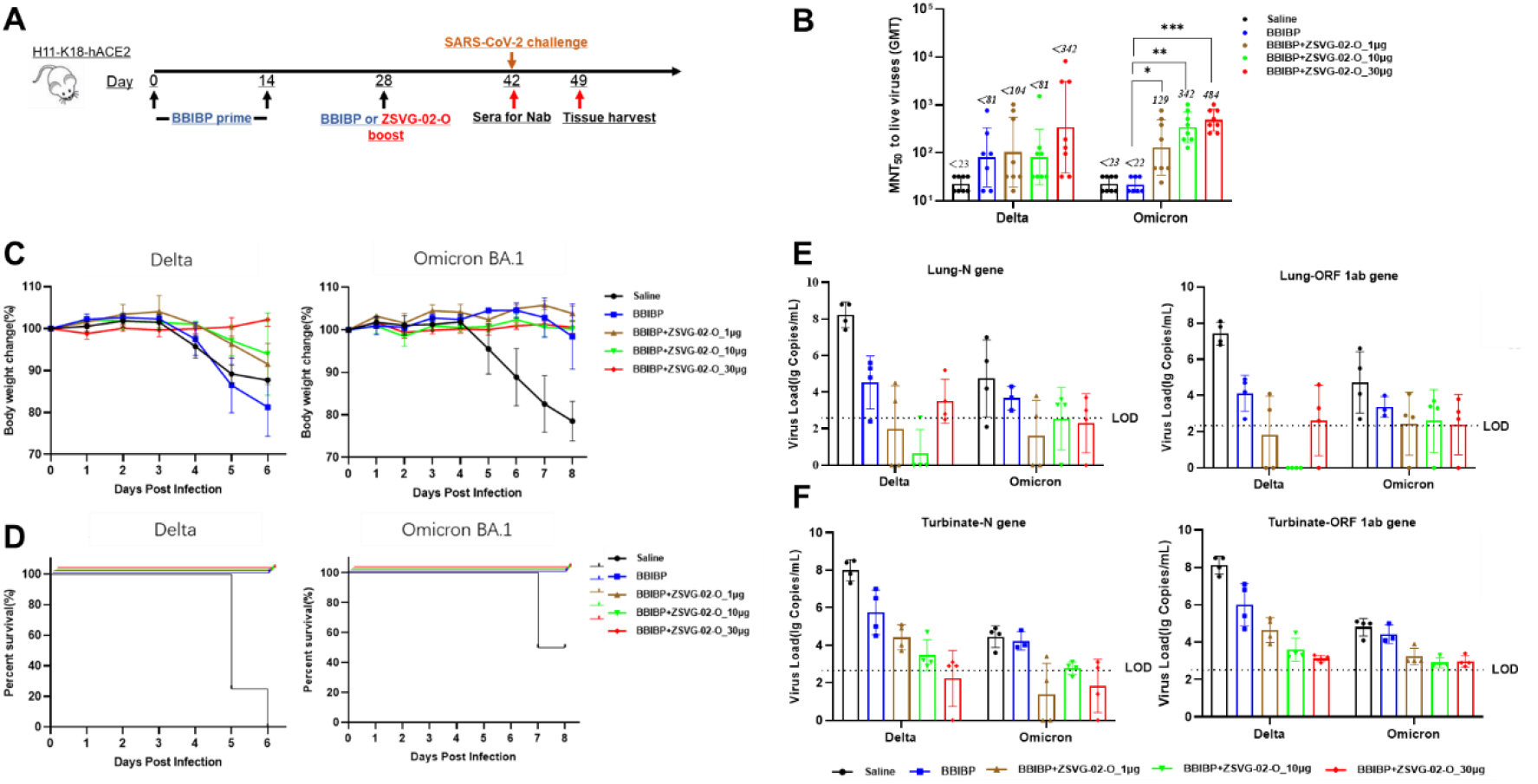
Protection against challenges of SARS-CoV-2 variants in BBIBP-ZSVG-02-O heterologous prime-boosted mice. (A) Schematic diagram of immunization regimens. (B) GMT of neutralizing antibodies against Delta and Omicron BA.1 live viruses on the day of virus challenges. (C) Body weight changes following virus challenges. (D) the survival rates post viral challenge. (E), (F) qPCR detection of viral loads in lung tissue and turbinate. LOD: limit of Detection.

### Two-dose ZSVG-02-O heterologous boosts induced more Omicron-specific neutralizing antibodies

We further investigated immune responses of a two-dose heterologous boost on BBIBP-CorV-primed mice. The sera were collected 13 and 14 days following the 1^st^ and 2^nd^ boost, respectively. Figure 6B and C compare NAb levels after 1^st^ and 2^nd^ boost for the BA.1 and BA.4/5 omicron subvariants. For the BA.1 variant, the second boost of BBIBP-CorV or 1 µg, 10 µg, 30 µg ZSVG-02-O induced 5.3-, 13.6-, 16.8-, and 10.6-fold NAb increases, respectively. For the BA.4/5 variant, the increases were 2.9-, 3.2-, 5.4-, and 3.1-fold. Thus, a second boost can noticeably increase NAbs against Omicron subvariants, especially the BA.1 subvariant. Further, compared to homologous boost (four shots of BBIBP-CorV), heterologous boost with ZSVG-02-O (two BBIBP-CorV + one or two ZSVG-02-O) induced much higher NAb levels for Omicron variants (Figure 6B-C). For BA.1, a single ZSVG-02-O boost induced NAb levels 2.3- to 7.1-fold and a double boost further increased the difference to 5.9- to 12.7-fold depending on the mRNA dose. For BA.4/5, the gap reached 1.3- to 9.6-fold after a single boost and 1.4- to 10.3-fold after a second boost.

**Figure 6:**
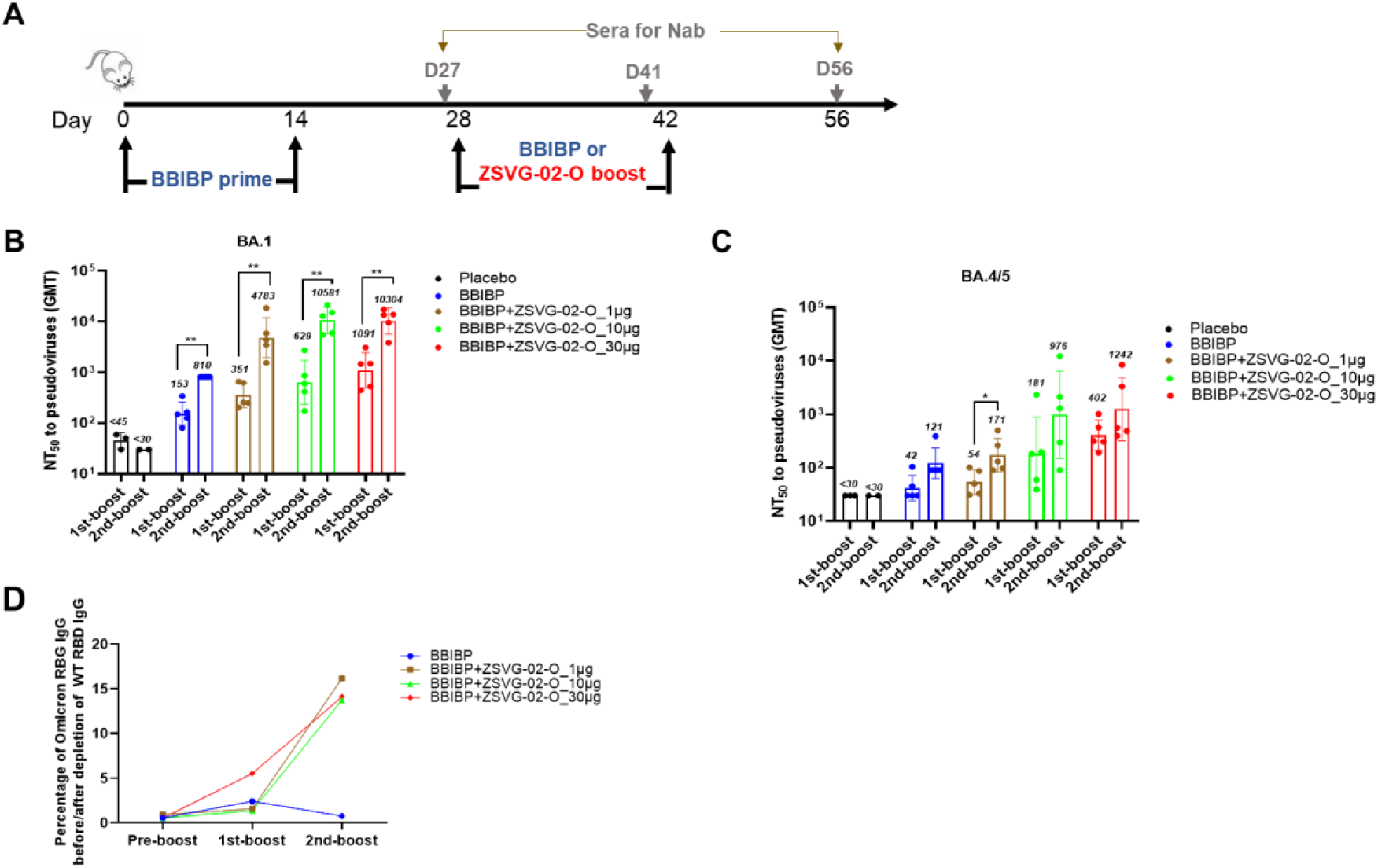
ZSVG-02-O boost induced Omicron-specific antibodies. (A) Schematic diagram of immunization regimens. Sera samples were collected at D27 (pre-boost), D41 (1st-boost) and D56 (2nd-boost). (B) GMT of neutralizing antibodies against Omicron BA.1. (C) GMT of neutralizing antibodies against Omicron BA.4/5. (D) GMT of Omicron RBD specific IgG after depletion of ancestral RBD-binding IgG.

Since one ZSVG-02-O boost to BBIBP-CorV-primed mice produces similar levels of NAbs for all three VOCs (Figure 4B and C), we wondered what population of antibodies, Omicron-specific or cross-reactive, this heterologous boost scheme produces. To investigate, we collected sera at Day 27 (pre-boost), Day 41 (13 days post 1st-boost) and Day 56 (14 days post 2nd-boost), and depleted their antibodies that react to the ancestral RBD. The resulting IgG was titrated with Omicron BA.1 RBD to determine the percentages of IgG for non-ancestral RBD. Our results show that after a single ZSVG-02-O boost, depletion of the ancestral RBD-binding IgG resulted in 95 - 99% loss of total IgGs in sera collected (Figure 6D). However, 2 weeks after the second boost with ZSVG-02-O, percentages of non-ancestral RBD-binding IgG increased to ∼14%, 3- to 10-fold that of a single ZSVG-02-O boost. As a control, we found no change in the percentages of non-ancestral RBD-binding IgG for animals receiving two doses of homologous BBIBP-CorV boosts. These results indicate that a single ZSVG-02-O boost following the BBIBP-CorV prime mainly elevates levels of antibodies cross-reactive to both Omicron and the ancestral RBD, while the second ZSVG-02-O boost specifically increases the population of antibodies that recognize Omicron RBD but ignore the ancestral RBD.

## DISCUSSION

The highly transmissible Omicron variants represent the ongoing challenge of battling the pandemic globally. Our data reveal that the ZSVG-02-O codes a highly dynamic Omicron Spike antigen that frequents multiple conformations, unlike the ZSVG-02 coded Delta Spike antigen. Our result, alongside most other recent studies, explains why mutations in Omicron RBD facilitate its binding to ACE2, and hence spreads more rapidly than Delta ^31, 32^. Our EM data also confirm that Omicron has evolved its Spike protein to adopt less stable thermodynamic states, presumably to facilitate evading host immune response while enhancing ACE2 interaction ^33^.

To counter these prevailing VOCs, this study introduces and evaluates the Delta-specific (ZSVG-02) and the Omicron-specific (ZSVG-02-O) mRNA vaccines. We demonstrate that they produce active antigen and induce a strong immune response in BALB/c mice with high neutralizing antibodies measured by live viruses. ZSVG-02 appears to induce higher neutralizing antibody levels for ancestral strain than ZSVG-02-O, probably due to the higher Spike protein homology between the Delta variant and the ancestral strain ^34, 35^. ZSVG-02-O likely produces more Omicron-specific antibodies that target Spike conformations otherwise unfavorable in Delta and ancestral virus. However, the Omicron BA.1-based ZSVG-02-O can produce antibodies that cross-react to Omicron BA.2 and BA.4/5 (Figure 2C), albeit with reduced activity due to progressive mutations in the two subvariants ^13, 36^. Nevertheless, our results show that ZSVG-02-O can induce sufficient levels of neutralizing cross reactivity against all Omicron subvariants, providing both humoral and T cell immunity.

ZSVG-02-O-stimulated humoral immunity differs from its T cell immunity. Our data show that ZSVG-02-O induces humoral response that skewed to their home variant in naïve animals (Figure 2). Conversely, T cell responses to the Spike proteins induced by ZSVG-02-O or ZSVG-02 appear similar across all three strains tested (Figure 3 and S3). The discrepancy of cross-reactivity between humoral and cellular immunity also emerges from people who received Ad26.CoV2.S or BNT162b2, or from recently convalescent but unvaccinated COVID-19 patients. Sera from those showed “diminished neutralization capacity” for Omicron variant but had similar T cell immunity across the ancestral, Beta, Delta and Omicron ^37^. T cell immunity provides protection through direct clearance of virus-infected cells by cytotoxic CD8+ T cells, or through enhancing humoral immunity by CD4+ T helper cells ^38^, without the need for humoral neutralizing antibodies. Clinically, higher levels of effector-associated transcripts in CD8+ T cell correlates with recovery from severe infections ^1^. The broad T cell immunity by vaccines may explain why ZSVG-02-O vaccination can protect animals from Delta virus challenges despite relatively lower NAb levels against the ancestral and Delta strains (Figure 2A).

We have investigated a heterologous prime-boost strategy with ZSVG-02-O following BBIBP-CorV, which is one of the major inactivated virus vaccines used in China and many other countries. From our data, we conclude that the humoral immunity induced by the heterologous BBIBP-CorV/ZSVG-02-O prime-boost regime differs greatly from that induced by two-dose ZSVG-02-O in naïve animals. When the naïve animals were vaccinated with ZSVG-02-O only, the resulting NAb levels clearly biases towards the Omicron variant (Figure 2). Conversely, a single ZSVG-02-O but not BBIBP-CorV boost following the two-dose BBIBP-CorV prime induces NAb levels comparable across all three strains (Figure 4). Our results on heterologous boost with ZSVG-02-O agree with recent studies ^39, 40^ on Omicron specific boosts. Those studies show a single Omicron-specific boost also induce similar antibody titers across the multiple strains. To understand the reason underlying the broadened cross-activity by the ZSVG-02-O boost, we analyzed the composition of total IgG in the sera from animals received the single dose ZSVG-02-O following the BBIBP-CorV prime. We found that the vast majority of antibodies induced by the first ZSVG-02-O boost were dual responders to both Omicron and ancestral RBDs. This result coincide with Gagne and coworkers’ report that, in their mRNA1273 primed monkeys, a single mRNA Omicron-specific boost only activated B cells that were dual responsive to both ancestral strain and Omicron while few Omicron-specific B cells were detected ^39^. Therefore, we postulated that a single ZSVG-02-O boost simply recalls the immunity built by the vaccine against the ancestral strain and “re-shapes” the IgG distribution to cross-react with Omicron, producing comparable NAb levels across all three strains tested. Thus, compared to two-dose immunization with Omicron-targeting vaccine alone, heterologous boost following vaccines for the ancestral strain can expand the breadth of immunity against more variants.

The immunity-broadening effect from BBIBP-CorV prime + ZSVG-02-O boost also manifested in enhanced T cell cross-reaction. In animals that received BBIBP-CorV vaccine followed by a single boost with the Delta-targeting ZSVG-02, the T cells show a strong response to the Nucleocapsid protein (N-protein) (Figure S5D) that is absent in naïve animals vaccinated with either ZSVG-02-O or ZSVG-02 alone (Figures 3 and S3). Thus, ZSVG-02 boost can recall existing cellular immunity generated by the BBIBP-CorV prime vaccination, since anti-N-protein immunity could only come from the inactivated whole-virus vaccine.

We further demonstrated that a second boost with ZSVG-02-O can significantly increase a population of Omicron-specific IgGs that do not cross-react with ancestral RBD (Figure 6D). This suggests that new B cell populations that produce Omicron-specific antibodies may necessitate a second Omicron-targeting boost. Alternatively, the Omicron-specific B cell population was already induced by the first boost, only needing additional time to produce sufficient levels of Omicron-specific antibodies. Resolving this mechanism warrants further NAb depletion assays using different and/or prolonged time points. In addition, further analysis of both germline and peripheral B-cell populations and their receptors responsive to S protein of SARS-CoV-2 variants is necessary to reveal the mechanisms. Meanwhile, we look forward to clinical studies that compare a ZSVG-02-O single- or double-boost in their abilities to protect against all current VOCs, particularly the Omicron variants.

Although homologous BBIBP-CorV prime-boost can partially protect the animals from Delta or Omicron BA.1 live viruses, we demonstrate that our heterologous immunization schemes offer complete protection against challenges by Delta and Omicron variants in a lethal ACE2-expressing mouse model. The BBIBP-CorV prime + ZSVG-02-O boost strategy produces higher NAb levels, broader T cell response, and hence more protection. It remains to be investigated whether a natural infection with Omicron variants in people previously vaccinated with the inactivated ancestral vaccines will produce similar immune responses as that by Omicron-targeting vaccine. In conclusion, our study proves that ZSVG-02-O effectively protects against Omicron infection as a standalone vaccine. More importantly, populations previously vaccinated with inactivated whole-virus of the ancestral strain, such as the BBIBP-CorV, can receive full benefits from a heterologous boost regime with ZSVG-02-O. The boost consolidates their previous primes and provides new resistance to Omicron variants by broadening both humoral and cellular immunity to cover all prevailing VOCs.

## MATERIALS AND METHODS

### Cryo-electron microscopy analysis

Both Spike proteins were expressed in HEK293F cells as previously reported ^41, 42^. The eluted proteins were concentrated using an Amicon Ultra centrifugal device (Millipore) with 100 kDa MW cut-off, and then applied to a Superose 6 5/150 column (Cytiva) for further purification. The peak fractions corresponded to S glycoprotein trimers were used for cryo-EM sample preparations. The proteins were concentrated to 0.75 mg/mL: An aliquot of 4 μL protein sample of Spike protein was applied onto a glow-discharged 300 mesh grid (Quantifoil Au R1.2/1.3) and GraFuture™-RGO grid which was supported with a thin layer of RGO (reduced graphene oxide), blotted with filter paper for 3.0 s using 3 blot force, and then plunge-frozen in liquid ethane using a Thermo Fisher Vitrobot Mark IV. Cryo-EM micrographs were collected on a 300 kV Thermo Fisher Titan Krios G4 electron microscope equipped with a Falcon4 direct detection camera. The micrographs were collected at a calibrated magnification of 96,000X, yielding a pixel size of 0.86 Å at a counting mode. In total, 5,248 micrographs were collected at an accumulated electron dose of 57.3 e-Å-2 s-1 on each micrograph that was fractionated into a stack of 32 frames with a defocus range of -1.0 to -2.0 μm. Beam-induced motion correction was performed on the stack of frames using MotionCorr2 ^43^. The contrast transfer function (CTF) parameters were determined by CTFFIND4 ^44^. A total of 5,248 good micrographs were selected for further data processing using the cryoSPARC software program according to standard procedures ^45^. Finally, the first conformation (2 up RBDs 1 down RBDs) was constructed from 432,771 particles and reconstructed at a global resolution of 2.82 Å based on the gold-standard Fourier shell correlation criterion at FSC=0.143 after refinement and CTF refinement. Another conformation (1 up RBDs 2 down RBDs) was executed by the same method, and the final resolution was 3.05 Å constructed from 346,652 particles. The molecular models were built by fitting a predicted structure of Omicron Spike protein from AlphaFold2 into the density map using UCSF Chimera ^46, 47^, followed by a manual model building of Omicron Spike protein in COOT ^48^ and real space refinement in PHENIX ^49^. The model statistics were listed in Supplementary Table 1.

### mRNA synthesis

mRNA was synthesized following standard procedures ^50-52^. Namely, we used T7 RNA polymerase in the presence of a CleanCap Reagent AG (3’ OMe) (TriLink) on linearized plasmids encoding codon-optimized SARS-CoV-2 gene. After transcription, the mRNA was purified by magnetic particles, flash frozen, and stored at -80°C until further use.

### mRNA Encapsulation

ZSVG-02 or ZSVG-02-O mRNA with four-component lipid mixture were fabricated as mRNA-LNPs using a microfluidic nanoprecipitation process in which an aqueous solution of mRNA was rapidly mixed with a solution of lipids. Next, the encapsulated mRNA was diluted to 0.5 mg/mL. Finally, the product was sterile filtered through a 0.22 µm Sartopore polyethersulfone (PES) membrane and aliquots were subsequently stored at room temperature or frozen at -80°C (1.0 mL fill).

### mRNA-LNP Characterization

Frozen vials were taken from storage and equilibrated to room temperature before characterizations. Particle size, polydispersity (PDI), and Zeta-potential of mRNA-LNPs were measured by dynamic light scattering (Malvern Nano ZS Zetasizer). Diameters are reported as Z-average. Encapsulation efficiency (E.E.) of mRNA in the LNP is defined as the mass ratio of encapsulated mRNA/total mRNA in the final mRNA-LNP product. Nucleic acids such as mRNA can be detected quantitatively using specific intercalating fluorescent dyes. The concentration of mRNA was calculated from a calibration curve generated using the mRNA standards. The relative proportion of encapsulated mRNA in mRNA-LNPs was determined via the ratio of the fluorescence signal in the absence vs. the presence of a surfactant that dispersed the LNPs. The signal in the absence of surfactant indicated the level of free mRNA while the signal in the presence of the surfactant provided a measure of the total mRNA in the sample.

To evaluate the quality and stability of the encapsulated mRNA, we compared ambient-stored LNPs with that after freeze/thaw cycles from -80°C. After freeze-thaw from -80°C, we observed almost identical size (∼90 nm) and monodispersity (PDI = 0.034) (Figure S1A) as the samples stored at room temperature (Figure S1B). We also measured similar encapsulation efficiency (E.E.) of 99.63% (ambient) and 98.02% (−80°C), showing both high purity and stability of our LNPs (Figure S1B). The slight negative Zeta potential of the mRNA-LNPs (−3 to -4 mV) can facilitate intracellular uptake (Figure S1C), as recommended by other common verified mRNA vaccine formulations ^53, 54^. Our LNP characterization results show that our procedure and formulation can produce monodispersed and stable mRNA-LNPs.

### Cell culture

HEK293T/17 cells were cultured in Dulbecco’s Modified Eagle’s Medium (DMEM, Gibco, USA) supplemented with 10% fetal bovine serum (FBS). The cells were maintained in a 37°C humidified incubator supplied with 5% CO_2_. The Vero cells (ATCC, TL-CCL-81.4) were cultured in DMEM (Gibco, USA) supplemented with 10% newborn calf serum (NCS, Gibco, USA), 50 U/mL penicillin-streptomycin (Gibco, USA), in a 37°C incubator with 5% CO_2_.

### Culture and Transfection of Human DCs

Frozen aliquots of human peripheral blood mononuclear cells (PBMCs) from healthy donors were purchased from Shanghai Saili Biotech. Six days before transfection, frozen PBMCs were thawed and resuspended in serum-free RMPI 1640 medium. Then PBMCs were plated in the cell culture dish with the density of 2 × 10^6^ cells/mL and incubated for 2 hours at 37°C. Next, the PBMCs were gently washed with warmed phosphate-buffered saline (PBS) three times. The adherent cells were cultured with RPMI 1640 medium supplemented with 10% FBS, human recombinant granulocyte-macrophage colony-stimulating factor (GM-CSF, 50 ng/mL R&D Systems, 7954-GM-020/CF), and interleukin-4 (IL-4, 50 ng/mL, R&D Systems, 6507-IL-025/CF) for 5 days. Then, the PBMC-derived immature dendritic cells (DCs) were replated into a 96-well plate. Cells were transfected with naked mRNA (500 ng/well), or 5′ppp-dsRNA (2500 ng/well; InvivoGen, tlrl-3prna), or 3p-hpRNA (Invivogen, tlrl-hprna-100), or PolyI:C (2500 ng/well; InvivoGen, tlrl-picw), or LNP-mRNA (500 ng/well) for 24 hours. The supernatant was collected for ELISA.

### Flow cytometry analysis

The HEK293T/17 cells were transfected with ZSVG-02 or ZSVG-02-O mRNA, either naked or LNP-encapsulated, at 0.75 µg/well. After 24 hours, the cells were harvested by 0.25% Trypsin-EDTA (Gibco) treatment and were blocked with 5% goat serum for 10 minutes. 100 µL biotinylated recombinant human ACE2 protein (His AVI Tag, 0.25 µg/µL, 1:100) (Sinobiology) and recombinant streptavidin protein (phycoerythrin) (Abcam) were used as the primary and secondary antibodies for ACE-2 protein detection. For RBD staining, PE-labeled Anti-SARS-CoV-2 Spike RBD neutralizing antibody (Acro) was used.

For T cell intracellular cytokine staining, splenocytes were isolated and resuspended with RPMI 1640 medium supplemented with 10% FBS. 2 μg/mL Spike trimer protein or peptide pool was added to media separately. Splenocytes were stimulated for 42 hours at 37°C before adding 2 mM monensin (Biolegend #420701) for an additional 6 hours. Then, splenocytes were collected by centrifuge and subjected to staining process. Firstly, cells were stained with live/dead cell dye (Zombie NIR™, Biolegend) as well as antibodies to cell surface markers (anti-45 AF700 [Biolegend, 147716], anti-CD3e FITC [Biolegend, 100306], anti-CD4 BV786 [Biolegend, 100552], anti-CD8 BV605 [Biolegend, 100744], anti-CD69 APC [Biolegend, 104514]) at 4°C for 30 minutes. Next, stained cells were washed with PBS, fixed and permeabilized using the Cytofix/Cytoperm™ kit according to manufacturer’s protocol (BD Bioscience, 554715). Finally, cells were stained with antibodies of cytokines (anti-IFN-gamma [Biolegend, 505808], anti-IL-2 [Biolegend, 503822], anti-IL-13 PE/Cy7 [Invitrogen, 25-7133-82]) at 4°C for 1 hour, and stored in PBS before analysis.

For effector cell and DC staining, uncultured splenocytes were incubated with antibodies (anit-CD45 APC/Cy7 [Biolegend, 103116], anti-CD11C FITC [Biolegend 117306], anti-CD3e Percp5.5 [BD Pharmingen, 551163], anti-IA/IE AF700 [Biolegend 107622], anti-CD4B V786 [Biolegend, 100551], anti-CD8 BV605 [Biolegend, 100744], anti-CD44 PE [Biolegend, 103008], anti-CD62L BV650 [Biolegend, 104453], anti-CD69 APC [Biolegend, 104514]) at 4°C for 30 minutes. The cells were then washed with PBS and subjected to flow cytometry analysis (Beckman, CytoFLEXLX).

### Enzyme-linked immunosorbent assay (ELISA)

To measure the secreted cytokine levels of IFNα and TNFα, the dendritic cell supernatant was collected at 24 hours post transfection with ZSVG-02 or ZSVG-02-O. Human TNF-α Ultrasensitive ELISA Kit (Invitrogen,KHC3014 [96 tests] and Human IFNα ELISA Kit [Invitrogen,BMS216TEN]) were used for cytokine quantification.

### Determination of viral titers by CCID_50_ assay

Virus-containing samples, Wild-type/Prototype/WIV04 (NCBI Reference Sequence: NC_045512.2), Delta (Genebank accession no. OU428631.1) and Omicron (GISAID accession no. EPI_ISL_7138045) were 10-fold serially diluted in infection medium DMEM + 2.5% NCS + 1% penicillin-streptomycin and then transferred into a 96-well plate with approximately 2 - 3 × 10^5^ Vero cells in each well with 8 replicates per sample. After 4 days of culture in a 37°C incubator with 5% CO_2_, cytopathic effect (CPE) was observed for the quantification of the median tissue culture infectious dose. The virus titer was calculated by the Karber method ^55^.

### Cell lysate preparation and Western blotting

The 293T/17 cells were transfected with ZSVG-02 or ZSVG-02-O mRNA produced by in vitro transcription (IVT), mock (empty lipofectamine 3000) and pcDNA3.1(-) plasmid-expressing S protein, respectively. After 24 hours, the cells were collected and lysed with RIPA buffer (Beyotime, P0013B). Equal amounts of protein samples were then separated by 10% sodium dodecyl sulfate-polyacrylamide gel electrophoresis (SDS-PAGE) followed by transfer to polyvinylidene fluoride (PVDF) membranes (Millipore, Billerica, MA, USA). The membranes were first blocked with 5% skim milk for 1 hour and then incubated with the primary antibodies of SARS-CoV-2(2019-nCoV)Spike RBD Antibody (Sinobiology, 40592-T62-100) for 2 hours at 37°C. After TBS-T (0.1% v/v) washing and incubation with the secondary antibodies of goat anti-rabbit IgG-Fc (HRP) (Sinobiology, SSA003), the protein bands were detected by LumiQ HRP Substrate solution (Share-bio, SSA003) and imaged with ChemiDoc™MP imaging system (Bio-Rad, USA)

### Pseudovirus Neutralization assay

The pseudovirus neutralization assays were performed according to protocols described previously ^1^. Briefly, serially diluted mouse serum samples were mixed with SARS-CoV-2 pseudoviruses (Vazyme Biotech Cat.# DD1402, DD1458, DD1768) in a 96-well plate and incubated at 37°C for 1 hour. Next, HEK293-ACE2 cells (Vazyme Biotech Cat.# DD1401) were added to each pseudovirus-containing well and incubated at 37°C for 48 hours. After incubation, the supernatants were removed and luciferase assay was performed with Bio-Lite™ Luciferase assay reagents (Vazyme Biotech, Cat.#DD1201). The neutralization antibody (NAb) titer was calculated and presented as half maximal effective concentration (EC50).

### Neutralization assay

Serum samples were diluted 1 in 16 in DMEM supplemented with 2.5% NCS, 50 U/mL penicillin-streptomycin (Gibco, USA), and then were added into an equal volume(0.05 mL/well)of challenge virus solution (100 CCID_50_ virus). After neutralization in a 37°C incubator for 2 hours, cell suspension with the virus was added into the 96-well plates (0.1 mL/well), which were cultured in a CO_2_ incubator at 37°C for 3 - 5 days. The assay of each serum was performed in 2 replicates, and the Karber method ^55^ was used to calculate the neutralization endpoint.

### Mouse Vaccination

All experiments were approved by Jiangsu Provincial Department of Science and Technology with an approval number of SYXK (Su) 2020-0041. BALB/c mice of 8 weeks of age purchased from China Jiangsu Huachuang Sino were immunized intramuscularly (IM) with ZSVG-02 or ZSVG-02-O on days indicated. Blood samples were collected for neutralizing antibody (NAb) detection and mice were harvested for T cell immunity experiments. For pre-immunization with BBIBP-CorV (0.5 mL, 6.5 U each tube), 100 µL BBIBP-CorV/mouse was IM-administered at Day 1 and Day 14, followed by ZSVG-02 or ZSVG-02-O boosting at Day 28. Mice were harvested at Day 42 for T cell immunity experiments.

### Mouse challenge with live SARS-CoV-2 Delta and Omicron BA.1 variant

The effect of *in vivo* protection by ZSVG-02 or ZSVG-02-O vaccination were assessed by using a *H11-K18-hACE2* mouse model with C57BL/6JGpt genetic background purchased from China GemPharmatec (Cat.# T037657) ^1, 56-59^. Briefly, groups of 3- to 4-month-old male mice were anaesthetized with isoflurane, followed by intranasal inoculation with 200 CCID_50_ of Delta or Omicron variant. Survival and body weight of the mice were monitored daily, and all animals were sacrificed upon death or at 6 or 8 dpi for harvesting lung tissues for virological assessment, pathological and histological examinations. Turbinate and lung tissues were subjected to qPCR viral load detection. The limit of detection (LOD) was approximately 2.7 Log copies/mL.

### Removal of ancestral strain RBD-binding IgG

The RBD-binding antibodies against the ancestral strain variant were depleted by using magnetic beads coated with the RBD fragment of ancestral strain SARS-CoV-2 (AcroBiosystems, MBS-K002). According to the manufacturer’s protocol, the beads were resuspended in ultrapure water at 1 mg beads/mL then washed 2 times in PBS with 0.05% BSA. The vaccine-immunized sera were diluted 8 times with 10 μL serum + 70 μL 0.05% BSA/PBS based on the manufacturer-reported binding capacity of 40 μg anti-RBD antibodies per mg beads. 80 μL beads were added to a tube and placed on the magnetic separator (Vazyme, CM101) for 1 - 2 minutes to remove the supernatant. Then, 80 μL of diluted serum was added to the tube and rotated at 4°C for 24 hours. The antibodies that bound RBD from the supernatant were separated by the magnetic separator, the supernatant was collected for another 80 μL beads for the second round of depletion to ensure full depletion of ancestral strain RBD-binding antibodies. The supernatants following the second depletion were collected and titrated for IgG that bind to Omicron RBD as described below.

### Serum antibody measurement by ELISA

The IgG ELISAs for RBD were conducted by using Antibody IgG Titer Serologic Assay Kit (AcroBiosystems, RAS-T059). According to the manufacturer’s protocol, the microplate of the kit was pre-coated with SARS-CoV-2 Spike RBD protein. Two-fold serial dilutions of sera were performed (serum dilution of D27 beginning at 1 in 25, serum dilution of D41 and D56 beginning at 1 in 250) in dilution buffer. Saline group serum was used as negative control and diluted at 1 in 100. 100 μL of diluted samples were added to the microplate and incubated at 37°C for 1 hour. After 3 washes, 100 μL HRP-goat anti-mouse IgG secondary antibodies were added and incubate at 37°C for 1 hour. After an additional 5 washes, 100 μL substrate solution were added to each well and incubate at 37°C for 20 minutes in the dark. Next, 50 μL of stop solution was added to each well. OD450 and OD630 were read on a Tecan Spark plate reader. The cut-off value was 0.1. The value of OD450-OD630 is greater than or equal to cut-off value is the positive reading, otherwise it is the negative reading. The antibody titer of the sample corresponds to the highest dilution factor that still yields a positive reading.

## ACKNOWLEDGMENTS

We thank Dr. Linglei Jiang for proofreading the manuscript. We thank Beijing institute of biological products Co., LTD. for providing BBIBP-CORV vaccine.

## Author contributions

WJ and XMY. designed and funded the research; XHC and WJ analyzed data; CL, YTZ, ZJW, ZBY, FJH, FL, GHY, RJC, LXL, JX, XW, KL, JD, XGL, QZ, CXH, P.P, RZ performed research; CL, YTZ, ZBY, FL, JX, GHY, RJC, ZJW and WJ prepared the manuscript; the rest of authors participated and performed the research.

