## supplemental table 1 for "Heterologous boost with mRNA vaccines against SARS-CoV-2 Delta/Omicron variants following an inactivated whole-virus vaccine"

**Table S1: Cryo-EM data collection, refinement and validation statistics**

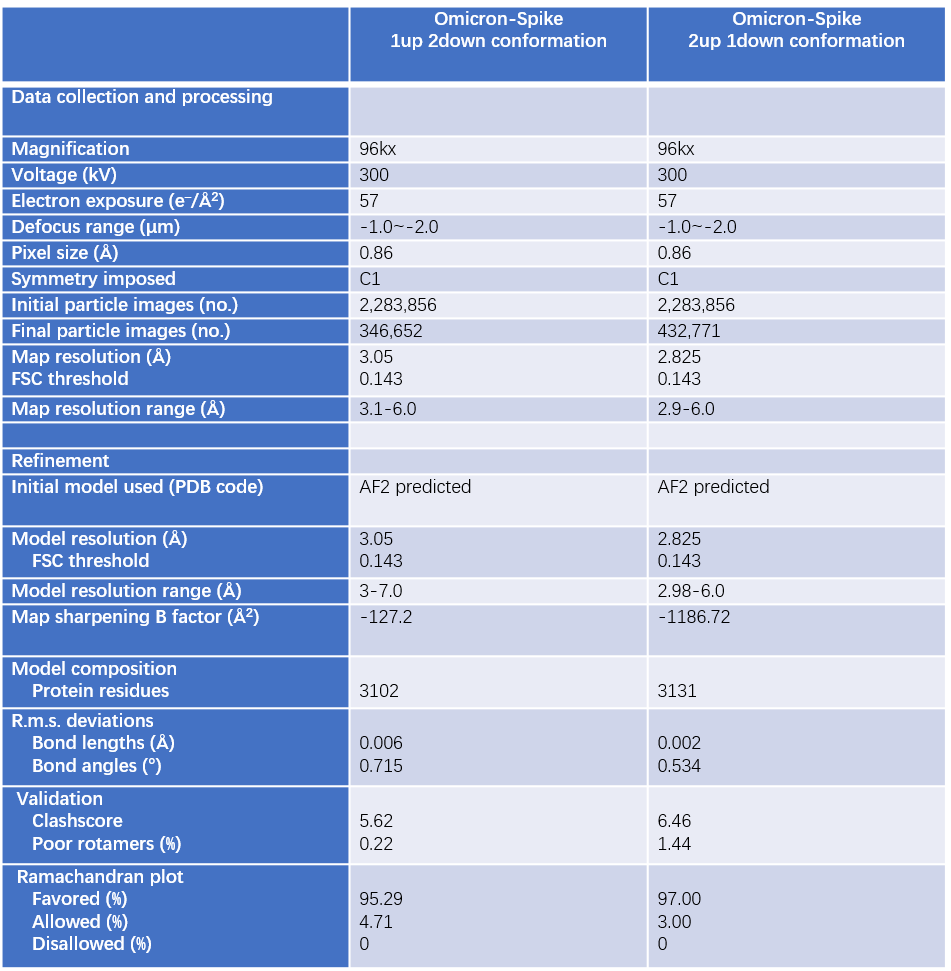

**Figure S1 Chareteritics of encapsulated mRNA**

(A) Representative TEM image of ZSVG-02-O mRNA-LNPs; B-D, Comparisons of mRNA-LNPs stored at room temperature and after freeze-thaw from -80oC, in terms of particle size(B), encapsulation efficiency (E.E.)(C) and Zeta potential (D).

(E). Quantification of S protein expression intensity from B, n = 4, independent experiments with ≥ 10000 cells each. MFI, mean fluorescent intensity, ***p < 0.0005, two-tailed ratio paired t-test.

(F), Inhibition of ACE2 binding to S-protein expressing cells transfected with ZSVG-02 by an anti-SARS-CoV-2 Spike RBD neutralizing antibody at different concentrations.

G, H Secreted cytokine levels of IFNα (G) and TNFα(H) 24 h after the transfection into dendritic cells, n = 3 independent experiments. Mock (empty lipofectamine 3000) as negative control, 5’ppp-dsRNA, 3p-hpRNA, poly(I:C) and Resiquimod, each of which stimulating different intracellular innate immune signaling pathways, as the positive controls.

(I) side view and (J) top view of different Delta and Omicron EM structures. The Delta spike active state with 1 up RBD is colored with corn silk.

(K) The Local resolution estimation result of each state EM map. Scales (Angstroms) to the right.

(L) The FSC curves of each state 3D reconstructions. The resolution values labeled near the FSC dash line is based on the gold-standard Fourier shell correlation criterion at FSC=0.143.

(M) The represented 2D classes results of Delta Spike (left) and Omicron Spike (right)

(N) The RMSD results of each Model chain compared with the references model indicated in the table.

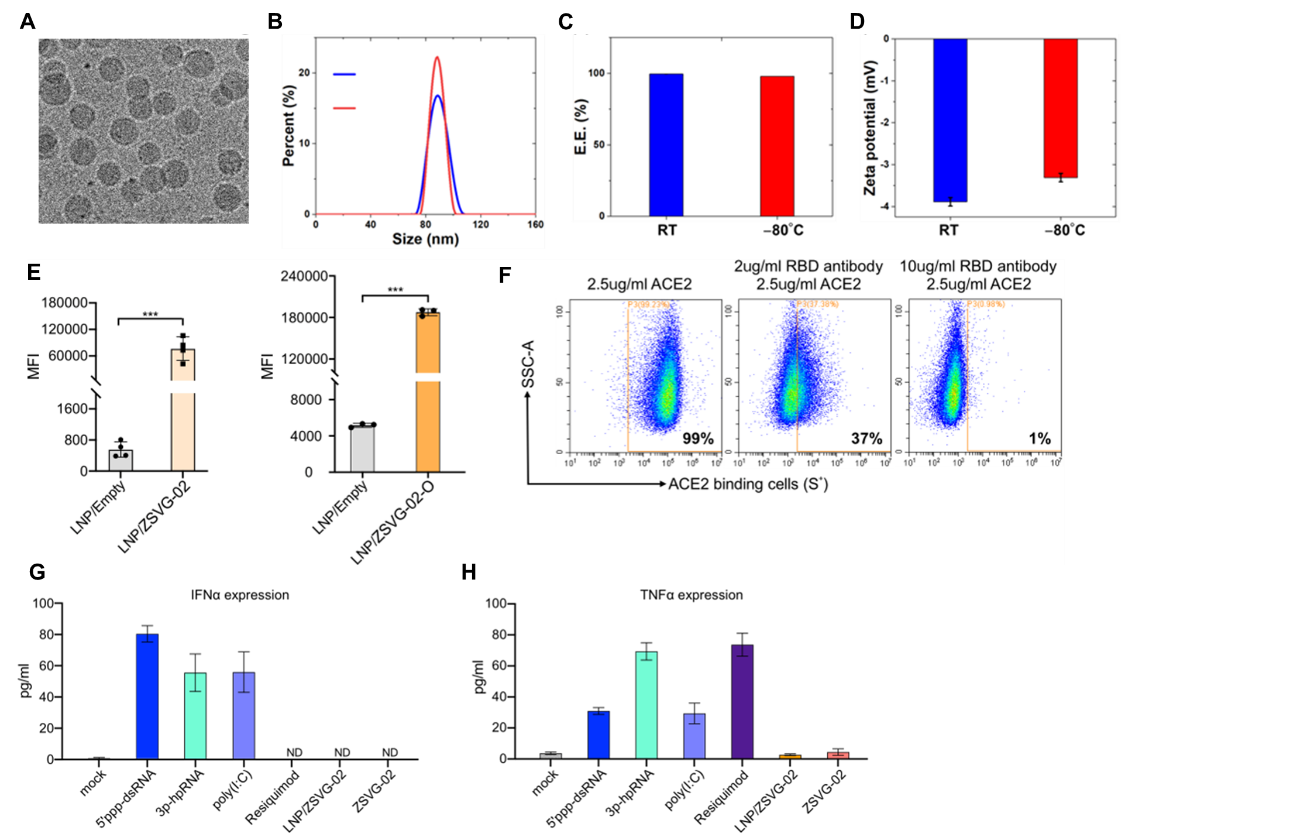

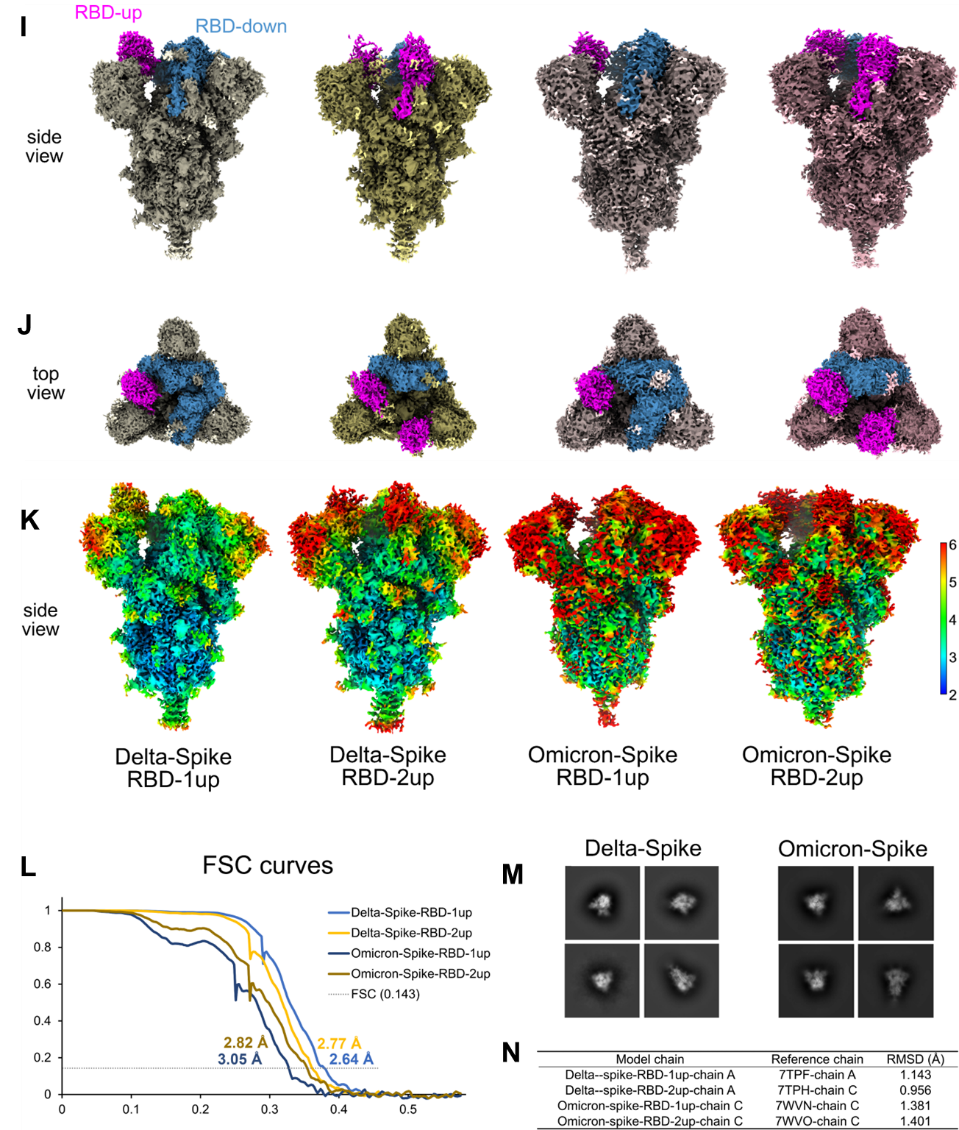

**Figure S2 ZSVG-02 induced antibodies cross-reacted with Omicron variants.** A, Schematic diagram of ZSVG-02 immunization and sample collection; B, GMT of neutralizing titers to pseudoviruses were determined 21 days post initial immunization; C, GMT of neutralizing titers to live viruses were measured 28 days post initial immunization. WT: ancestral strain

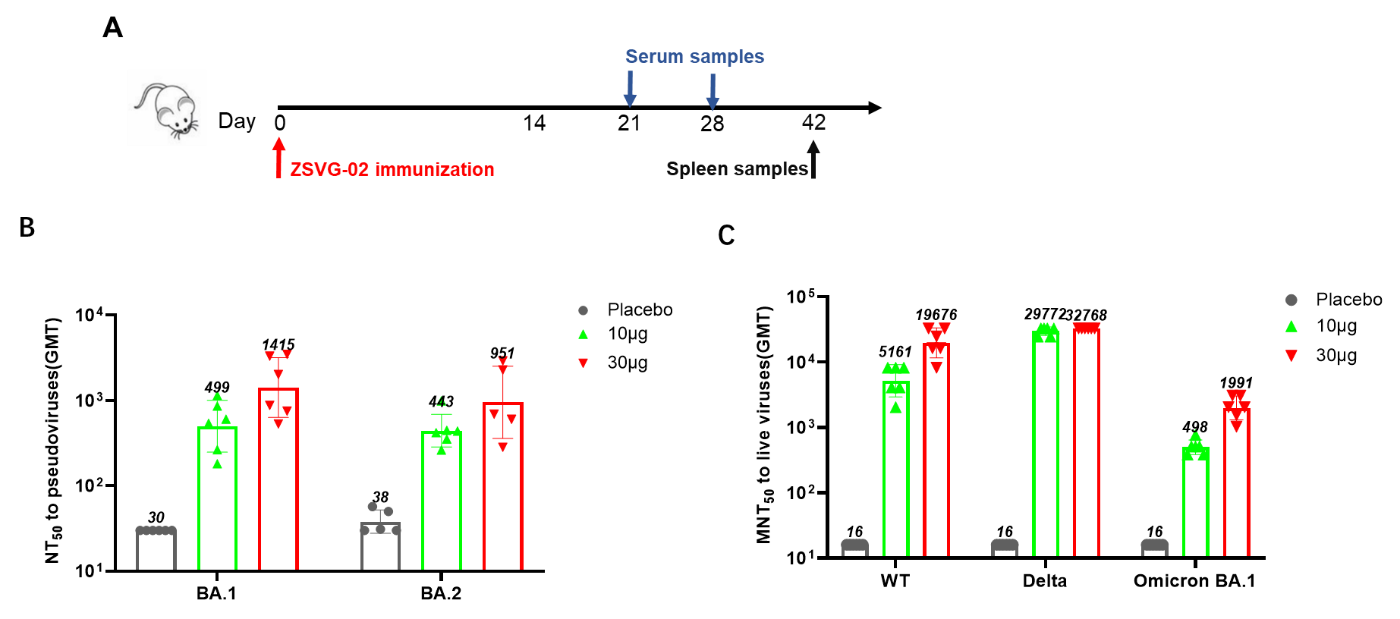

**Figure S3. ZSVG-02 immunization induced T cell immunity**

Splenocytes were collected 4 weeks post second immunization and stimulated with the indicated spike protein of different variants for 48 hours. A-H,The proportions of IFNγ+/CD69+ T cell in CD4 and CD8 T cells determined with flow cytometry after restimulation; I, IL13 expression in CD4 T cells determined with flow cytometry after restimulation; J-K, ELISPOT assay for IL2 and IL-5 in splenocytes stimulated with delta+ specific peptides. WT, ancestral strain.

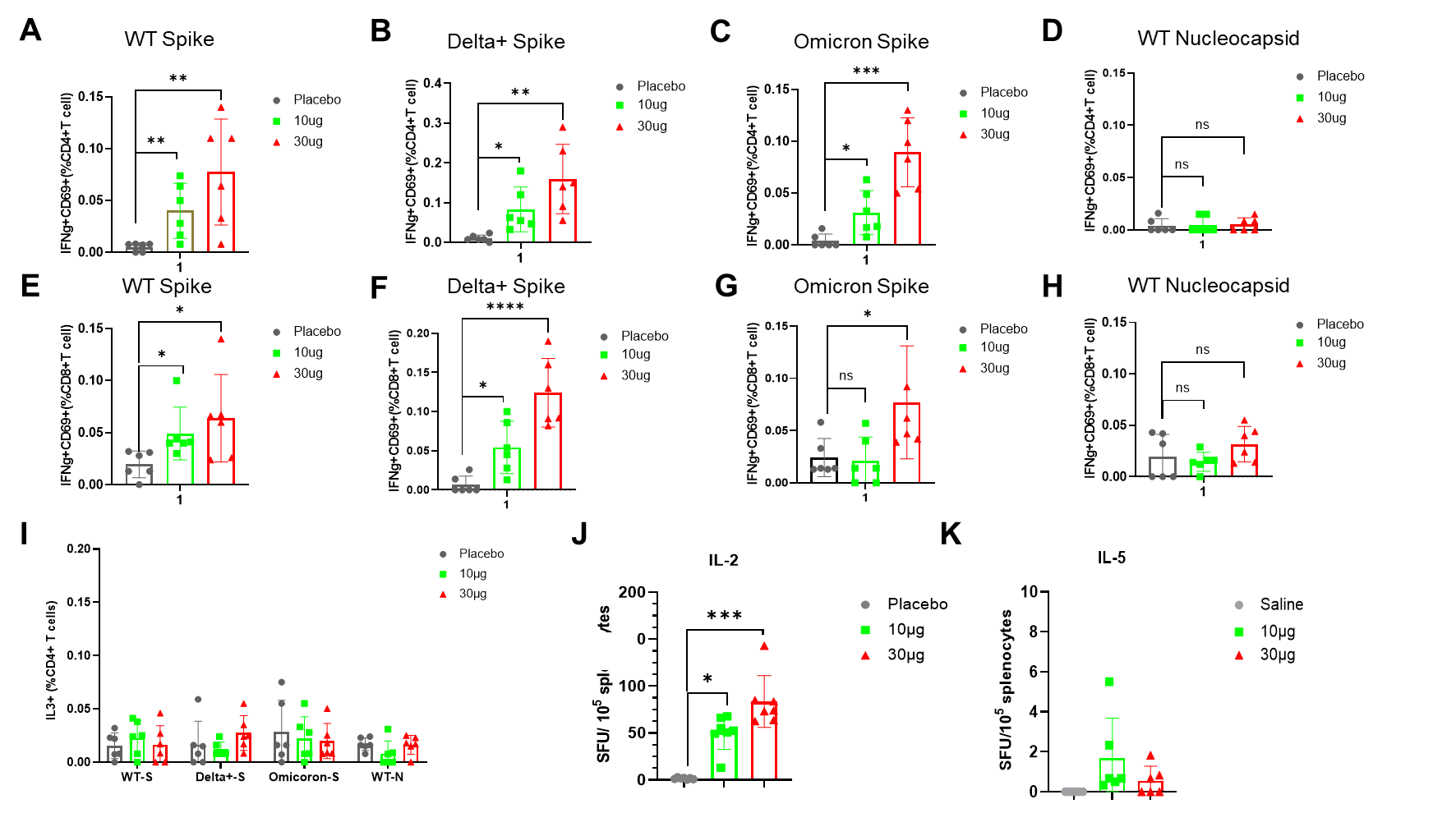

**Figure S4. Serological evaluation and T cell immunity in mice with ZSVG-02 boost following two doses of inactivated SARS-Covid-2 vaccine.**

A, Schematic diagram of immunization regimens for ZSVG-02 boost; B, Neutralizing antibody titers were measured with pseudovirus neutralization assay in mice boosted with ZSVG-02; C-H, Splenocytes were collected 4 weeks post second immunization and stimulated with the indicated spike protein of different variants including ancestral, Delta, Omicron and C.1.2 variants for 48 hours. The proportions of CD69+ T cell in CD4 and CD8 T cells determined with flow cytometry after restimulation; I-J, The proportions of Tfh (Foxp3-PD1+) and DC (CD11C+IA/IE+) in spleen. A single dose boost with ZSVG-02. WT, ancestral strain.

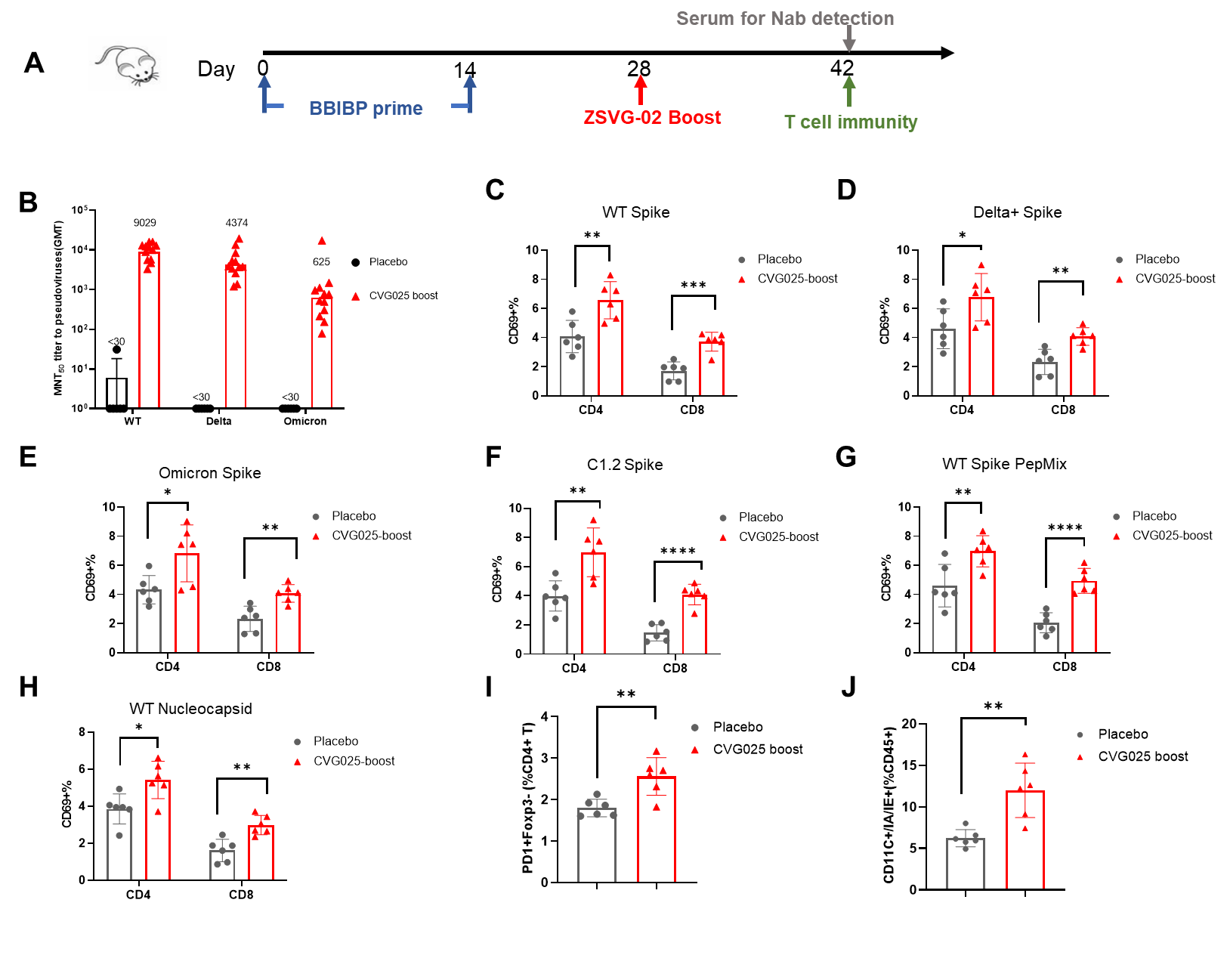

**Figure S5: Protection against SARS-CoV-2 Delta and Omicron variants challenge in ZSVG-02 vaccinated mice.** A, Schematic diagram of immunization regimens. B, Body weight changes in each group post infection with Delta variant (Left, B.1.617.2 strain) and Omicron variant (Right, B.1.1.529 strain). C, the survival rates post viral challenge. D-E, qPCR detection of viral loads in lung tissue and turbinate. LOD: limit of Detection. Data are shown as mean ± SEM (****p < 0.0001).

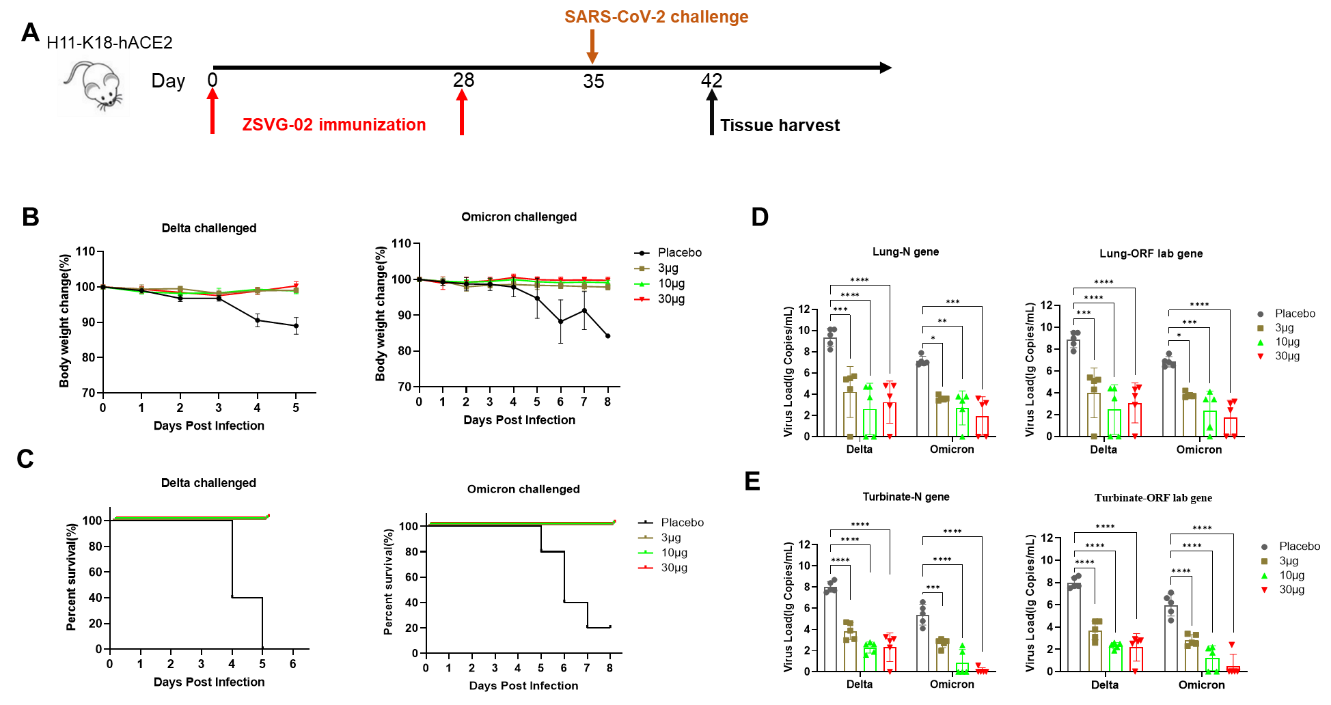
